# Interaction of the Motor Protein SecA and the Bacterial Protein Translocation Channel SecYEG in the Absence of ATP

**DOI:** 10.1101/799247

**Authors:** Klemens Winkler, Andreas Karner, Andreas Horner, Christof Hannesschlaeger, Denis Knyazev, Christine Siligan, Mirjam Zimmermann, Roland Kuttner, Peter Pohl, Johannes Preiner

## Abstract

Translocation of many secretory proteins through the bacterial plasma membrane is facilitated by a complex of the SecYEG channel with the motor protein SecA. The ATP-free complex is unstable in detergent, raising the question how SecA may perform several rounds of ATP hydrolysis without being released from the membrane embedded SecYEG. Here we show that dual recognition of (i) SecYEG and (ii) vicinal acidic lipids confers an apparent nanomolar affinity. High-speed atomic force microscopy visualizes the complexes between monomeric SecA and SecYEG as being stable for tens of seconds. These long-lasting events and complementary shorter ones both give rise to single ion channel openings of equal duration. Furthermore, luminescence resonance energy transfer reveals two conformations of the SecYEG-SecA complex that differ in the protrusion depth of SecA’s two-helix finger into SecYEG’s aqueous channel. Such movement of the finger is in line with the power stroke mechanism of protein translocation.

## Introduction

An important step in the biogenesis of many secretory proteins is their translocation across the lipid bilayer. The conserved Sec protein translocation machinery facilitates most of these translocation events. In bacteria it contains two key elements, the hetero-trimeric transmembrane channel SecYEG and the soluble motor protein SecA. While SecYEG provides the pathway for protein translocation, SecA affords part of the translocation energy by hydrolyzing ATP, the other part being provided by the proton motive force.^1^ SecA gains access to the SecYEG complex via a lipid-bound intermediate state.^2^ Its highly amphipathic N-terminal helix carries positively charged amino acids aligned on one side, and hydrophobic amino acids on the other which interact with negatively charged phospholipids.^3^ The N-terminus thereby penetrates into the membrane and aligns parallel to the membrane-plane at a depth of 7-8 L.^4^ Deletion of the first 20 amino acids of the N-terminus (SecA His-ΔN20) impedes SecA dimerization.^5, 6^ The accompanying activity loss can be reversed by substituting the N-terminus for a His-tag and supplementing SecYEG proteoliposomes with Ni^+^ - NTA lipids, suggesting that membrane tethering is important for functioning.^7^ However, the crystal structure of the SecA-SecYEG complex^8^ indicates that the helical part of the amphipathic SecA N-terminus is not in contact with the membrane. Thus, a major conformational change of SecA was suggested to allow SecYEG binding and penetration of the N-terminus at the same time.^2, 4^ Yet, structural evidence for SecA’s large conformational change is lacking.

The above-mentioned experimental observations have been made with SecA bound to nucleotides. However, during translocation of a polypeptide, each nucleotide remains bound to SecA only for a very limited time.^9^ A single ATP hydrolysis event energizes the movement of only a few amino acids. Energizing the movement of the next protein segment requires ADP release and the binding of a new ATP molecule, which includes a timespan during which SecA is effectively in a nucleotide-free conformation. The affinity of the nucleotide-free conformation to the lipid membrane is not known. It has only been reported that nucleotide release weakens the interaction between SecA and SecYEG^10^, and as a consequence, the SecA SecYEG complex appears to be unstable in detergent.^8^ It is thus unclear how SecA may perform many hydrolysis cycles^7^ without being released from SecYEG.

Using a set of complementary biophysical techniques, we were able to reconstruct the detailed molecular pathway of SecA targeting SecYEG embedded in a lipid membrane and unravel how SecA stays bound to SecYEG during the time-interval between ADP release and ATP binding. We built a coherent mechanistic model based on kinetic rate constants and structural information obtained from surface plasmon resonance (SPR) experiments, high-speed atomic force microscopy (HS-AFM)^11–14^, luminescence resonance energy transfer (LRET),^15, 16^ and single channel electrophysiological recordings.^17–20^

## Materials and Methods

### SecA purification and labeling

SecA was obtained from overexpressing E.coli cells as previously described.^18^(Knyazev et al., 2020) To position an acceptor dye on the tip of SecA’s two-helix finger, we introduced a cysteine at position K797 by site directed mutagenesis into the expression vector pET30b SecA(N95). This mutant was previously shown to have minimal influence on the translocation activity of SecA.^21–23^ After induction at an O.D. 600 of 0.9 with 1mM Isopropyl-β-D-thiogalactopyranosid (IPTG, Peqlab, VWR) SecA(N95) K797C was overexpressed for 4 hours at 37 °C in E.coli NiCo21 cells (New England Biolabs) grown in 2xYT (Acumedia) containing Kanamycin. Cells were pelleted and lysed using an Avestin Emulsiflex C5 in Lysis Buffer (20 mM HEPES pH 7.5, 0.5 M NaCl, 25 mM Imidazole, supplemented with Protease inhibitors) with 2 cycles of 20,000 psi. After 1 hour of centrifugation at 100,000xg at 4 °C the supernatant was incubated with Chitin resin (New England Biolabs) for 30 minutes at 4 °C to remove metalloproteases. SecA(N95) K797C was affinity purified by incubating the flow-through with Ni^2+^-chelating beads for 1 h at 4 °C. Columns were washed with Wash Buffer (20 mM HEPES pH 7.5, 0,5 M NaCl) containing 25 mM imidazole. Beads were washed with Wash Buffer before incubation with 0,6 mM TCEP for 10 minutes. Labeling was performed using Atto488-maleimide (Atto-Tec) for 45 minutes. Following another washing step with Wash Buffer labeled SecA(N95) K797C was eluted in Elution Buffer (20mM HEPES pH 7.5, 0.5 M NaCl, 200 mM Imidazole) and concentrated to 500μl. Finally, we performed size exclusion chromatography on an ÄKTA pure system (GE Healthcare, Little Chalfont, UK) equipped with a Superdex 200 Increase 10/300 GL size exclusion column (GE Healthcare) using 20 mM HEPES pH 7.5, 100 mM NaCl. Protein concentrations were assessed using Bradford assays.

### SecYEG purification

SecYEG was purified and labeled as previously described.^19^ In short, SecYEG was obtained from E.coli BL21 (New England Biolabs) transformed with the arabinose dependent pBAD vector encoding cysteinless SecE, SecY, and SecG genes with an LBT tag either at G297 or at E62 of SecY. The G297 position is located in a loop of the periplasmic end of transmembrane helix 7. Since it is part of the lateral gate, it is likely to change position upon channel opening. The same holds true for E62 due to its location on SecY’s plug.

Transformed E.coli cells were grown in 2xYT medium (Acumedia) containing Ampicillin to an O.D. 600 of 0,7 at 37 °C when overexpression was induced by the addition of 2g/l arabinose. After additional incubation for 5 hours at 37 °C cells were harvested by centrifugation and resuspended in Lysis Buffer (20 mM Tris pH 7.5, 300 mM NaCl, 10% glycerol) supplemented with protease inhibitors (Thermo Scientific Halt Protease Inhibitor Cocktail and 0,5 mM AEBSF). After lysis with 2 cycles of 20,000 psi using an Avestin Emulsiflex C5 membrane fractions were separated by centrifugation at 100,000xg for 1 hour at 4° C and membrane pellets were solubilized in Lysis buffer containing 1% DDM (n-Dodecyl-β-D-maltopyranoside, Anatrace). Following incubation for 1 hour at 4° C solutions were centrifuged at 100,000xg for 30 minutes at 4° C. Supernatants were incubated with Ni-NTA resin (Ni-NTA Superflow, Qiagen) for 1 hour at 4° C and washed with Wash Buffer (20 mM Tris pH 7.5, 300 mM NaCl, 10% glycerol, 0,6 mM DDM, 20 mM imidazole). Proteins were eluted using 10ml of Wash Buffer containing 200 mM imidazole. The eluate was concentrated to 500μl and size exclusion chromatography was performed on an ÄKTA pure system (GE Healthcare, Little Chalfont, UK) equipped with a Superdex 200 Increase 10/300 GL size exclusion column (GE Healthcare), equilibrated with SecYEG FPLC Buffer (20 mM Tris pH 7.5, 100 mM NaCl, 10% glycerol, 0.6 mM DDM). Protein concentrations were assessed using Bradford assays.

### proOmpA purification

The pTrcHis proOmpA vector was transformed into temperature-sensitive E.coli MM52 cells. Cells were grown in 2xYT medium (Acumedia) containing Ampicillin at 30 °C until an O.D.600 of 0.9 was reached. After a 1:10 dilution with prewarmed 2xYT medium and further incubation at 37 °C for 30 minutes, over-expression was induced by the addition of 1 mM IPTG. After 2 hours cells were harvested. Cell lysis was performed in 50 mM Tris-HCl pH 8.0, 10% v/v glycerol, 300 mM KCl supplemented with protease inhibitors by 2 rounds at 20,000 psi in a homogenizer Emulsiflex C5 (Avestin). Insoluble material was harvested by ultracentrifugation at 4 °C and 100,000xg, for 25 minutes. After resuspension in Solubilization buffer (50 mM Tris-HCL pH 8.0, 6 M Urea) and stirring of the sample at 4 °C for 1 hour solubilized fractions were obtained by another ultracentrifugation step at 4 °C and 100,000xg for 90 minutes. Supernatant was incubated with Ni^2+^-chelating beads (Ni-NTA superflow, Qiagen) for 1 hour. After washing (50 mM Tris-HCl pH 8.0, 6 M urea, 10 mM imidazole) proOmpA was eluted with Elution buffer (50 mM Tris-HCl pH 8.0, 6 M urea, 300 mM imidazole) and dialysed over-night against 50 mM Tris-HCl pH 8.0, 6 M urea.

### SecYEG reconstitution into lipid vesicles and planar lipid bilayers

SecYEG complexes were reconstituted into lipid vesicles and planar bilayers as previously described.^17^ In brief, to 20 mg/ml of E. coli polar lipid extract (Avanti Polar Lipids, Alabaster, AL) we sequentially added 50 mM K-HEPES pH 7.5, 6% deoxy Big-CHAP (Affymetrix Anatrace, Maumee, OH, USA) and SecYEG in detergent (Protein to Lipid ratio of 1:50) at room temperature. For LRET experiments a DOPE:DOPG (Avanti Polar Lipids, Alabaster, AL) mixture (mass ratio 7:3) was used instead of E.coli polar lipid extract due to unfavorable unspecific binding of Tb^3+^ to this lipid extract. Subsequent to detergent removal by Bio-beads SM2 (Bio Rad), the proteoliposomes were harvested by ultracentrifugation (80 min at 100,000 x and resuspended at a lipid concentration of 5–10 mg/ml in a buffer containing 50 mM HEPES (pH 7.0), 10% glycerol and protease inhibitor. To show channel functionality we tested reconstituted SecYEG mutants for their translocation capability as previously described.^24,25^ In the presence of both ATP and SecA proOmpA-DHFR is translocated into proteo-LUVs and is therefore not accessible to a subsequent proteinase K digest (Fig. S1).

### Electrophysiology

Single channel measurements were performed as previously described.^18,19^ Ag/AgCl reference electrodes in the cis and trans compartments were to the command signal of the patch clamp amplifier (model EPC9, HEKA electronics, Germany) and the ground, respectively. The recording filter for the transmembrane current was a 4 pole Bessel with −3 dB corner frequency of 0.1 kHz. The raw data were analyzed using the TAC software package (Bruxton Corporation, Seattle, WA). Gaussian filters of 12 Hz or 112 Hz were applied to reduce noise.

To perform electrophysiological measurements we fused proteoliposomes to preformed planar bilayer lipid membranes. These “solvent-free” planar bilayers were folded by raising the level of two adjacent aqueous solutions over the dividing aperture in a Teflon septum with E. coli polar lipid extract (Avanti Polar Lipids, Alabaster, AL) monolayers on top.^26^ Fusion was facilitated by a 500 mM: 100 mM KCl gradient across the planar membrane. The hyperosmotic compartment contained the proteoliposomes and SecA. Both compartments were buffered by 25 mM K-HEPES at pH 7.5. In addition to the wild type translocon, we also subjected the mutants with LBT tag and fluorescent dye to our electrophysiological experiments. The result was invariant.

### LRET

Tb^3+^ acted as LRET donor in our experiments. It was embedded in a high affinity (K_d_ = 57±3 nM) lanthanide binding tag that consisted of the following 17 amino acids: YIDTNNDGWYEGDELLA.^27^ Its tryptophan served as antenna for efficient Tb^3+^excitation at 266 nm by a Nd:YAG-laser system (VM-TIM, Jena, Germany). We used a home-built LRET setup as described.^28^ Briefly, the laser light passed a dichroitic mirror (z266rdc, Chroma, Bellows Falls, VT, USA) and was focused on the sample via a 40 x 1.25 glycerol immersion quartz objective (Sysmex Partec GmbH, Münster, Germany). The objective also served to collect the light emitted by Tb^3+^ or the acceptor molecules, Atto488 or Atto532. These fluorescent dyes were attached via maleimide linkers to genetically introduced cysteines in SecA. Both Tb^3+^ luminescence and the fluorescent light further went through the dichroic mirror to a longpass filter (HQ465lp, Chroma, Bellows Falls, VT, USA), which removed residual excitation light. Close proximity between Tb^3+^ and the fluorescent dye resulted in a decrease of luminescence lifetime. Alternatively, to donor luminescence, we measured acceptor fluorescence (sensitized emission). For that purpose, a bandpass filter (D520/25m or HQ567/15x, Chroma, Bellows Falls, VT, USA) was additionally inserted into the emission beampath. In both measurement modes the light was absorbed by a gated single photon avalanche diode (SPCM-AQRH, Perkin Elmer, Waltham, MA, USA), connected to a DSN-101 power supply (PicoQuant, Berlin, Gremany). The signal was recorded and triggered by a multichannel scaler card (NanoHarp 250, PicoQuant, Berlin, Gremany). A delay of 30 μs between the onset of the light pulse (5-6 ns pulse duration) and diode activation helped to avoid any interference of the measurement with direct acceptor excitation.

As a system check, we used parts of the two-helix-finger, the C-terminal α-helix of SecA, as a molecular ruler. We measured a distance of 47 Å for the unbound SecA K797C-Atto488 V829LBT double mutant, which perfectly agrees with estimates from crystal structures (2fsi – 47.5 Å, 2fsh – 47.1 Å, 2ipc – 49.7 Å).

### Evaluation of luminescence decay curves

Even though several publications^16,29–31^ point out how LRET decay curves may be evaluated it is worth to lay out the basis for the global fit routine employed in our evaluation.

A given number of donors *D*_0_ is excited by a nanoseconds laser pulse, which appears on the timescale of the observed decay as instantaneous. Each donor either can decay directly or may – if present-transfer its energy to an acceptor. This occurs with a time constant *τ*_*T*_. Since the acceptors emit within nanoseconds, this process is considered as instantaneous as well. The population of donors may be split in several fractions where no energy transfer occurs (fractions *a*_*D*_ = *D*_*D*_/*D*_0_) or where transfer occurs with distinct time constants *τ*_*T,i*_ (fractions *a*_*i*_ = *D*_*i*_/*D*_0_). For the donors *D*_*D*_, the decay is simply described by

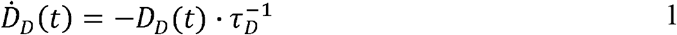

With the solution

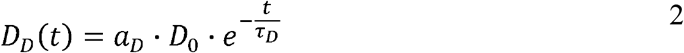

Since the detected counts (*counts*_*tot*_ (*t*)) are proportional (proportionality factor *α*,) to the number of emitted photons per time, equations 1 and 2 yield (*BG* is the detector background):

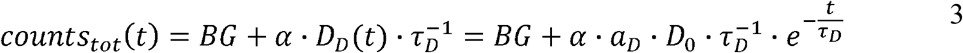

If LRET occurs, the following differential equation describes the two pathways that are possible (direct decay or energy transfer):

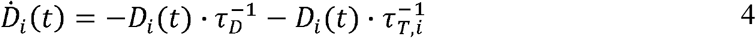

This is solved by:

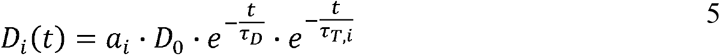

If sensitized emission is detected (by inserting a bandpass filter into the emission beam path), only the donors decaying into the LRET pathway are detected (the proportionality factor *α*′ and *BG*’ in this detection mode can differ from *α* and *BG* since the detection efficiency of the detector may be wavelength dependent or the altered acquisition settings are required for sensitized emission compared to total emission):

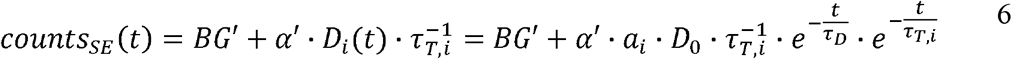

If the total emission is detected, one collects light from both decay pathways:

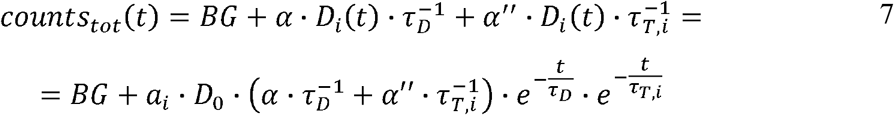

Again, a proportionality factor *α*″, is introduced since photons that are emitted directly are not detected with equal efficiency as compared when LRET has occurred.

We describe a system, where some donors have no acceptor present, some have one that sits close (short transfer time *τ*_*T,i*_), some have one that sits further away (long transfer time *τ*_*T,i*_).

Hence the sensitized emission becomes:

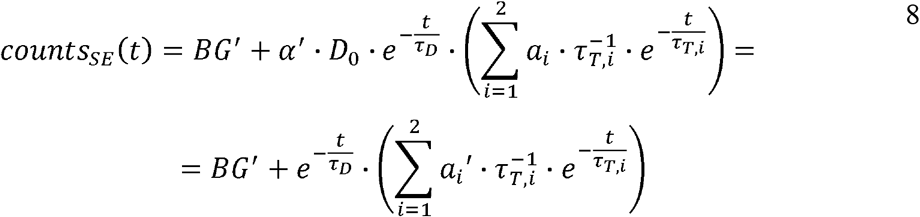

The last equity employing 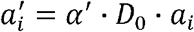 removes dependent variables that would hamper the fitting routine.

The total emission results in (combination of equation 3 and 7):

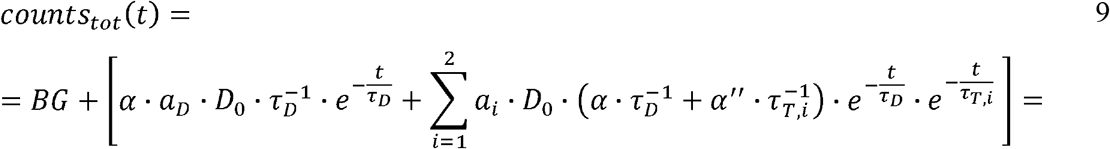

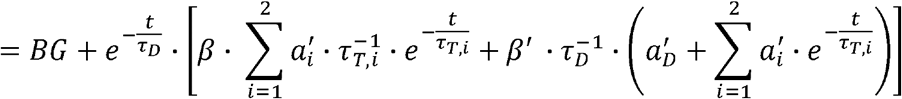

Again, the last equity employs 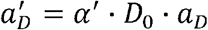, *β*′ = *α*/*α*′ and *β* = *α*″/*α*′ to remove dependent variables. The actual fractions are obtained via:

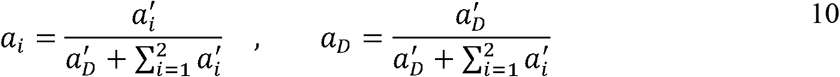

Recorded decay curves were fit with the NonlinearModelFit-Routine of Mathematica 11.2 (Wolfram Research, Champaign, USA). A weight of 1/y was applied in the routine. A titration experiment consisting of pairs of total emission and sensitized emission curves is globally fit in the following way:

- All traces share the same lifetimes of the donor (*τ*_*D*_) and transfer lifetimes *τ*_*T,i*_
- All traces share the same proportionality factors *β*′ and *β*
- Each pair of total emission (fit with equation 9) and sensitized emission (fit with equation 8) share the same 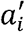 and 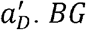 and *BG*′ are local parameters.
- If no acceptor is present in the system (e.g. 0 μM SecA in Fig. 4a and b), the parameters 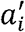 are set to 0 for this pair of traces.

Distances *r*_*i*_ between donor and acceptor are calculated via (*R*_0_ is the Foerster radius of the donor-acceptor pair)

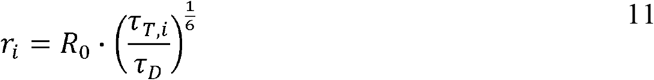

The distance between the C_α_ atoms of SecYEG G297LBT with SecA K797C is equal to approximately 46 Å for the PDB entries 5GAE, 5CH4, 5AWW, 5EUL, 3J46. Opening of lateral gate is predicted to decrease it to 43.9 Å (3J45).^32^ The maximal distance was equal to 50.3 Å for a conformation with a closed gate (3DIN).

From SecYEG E62LBT to SecA K797C the distance varies between 30 Å and 40 Å, depending on the position of the plug in the crystal structure. Both distance estimates are in the range of the Foerster radii of the donor-acceptor pairs (Tb^3+^ - ATTO 488, R_0_=39,72 Å and Tb^3+^ - ATTO532, R_0_=48,14 Å). These radii are calculated form the spectra of the donor and acceptor ^29^.

As ATP binds Tb^3+^ stronger than the lanthanide binding tag, LRET experiments were exclusively performed under nucleotide free conditions. Presence of proOmpA did not significantly change any of the observed characteristics.

### Fluorescence Correlation Spectroscopy (FCS)

To estimate the amount of liposomes and protein abundance in proteoliposomes, samples were subjected to fluorescence correlation spectroscopy (FCS) as previously described.^33^ After having counted the number of proteoliposomes, the samples were dissolved by detergent (1% Octyl glucoside, 2% Deoxy Big CHAP) and the newly formed micelles were counted again. By (i) assuming that every micelle contained exactly one SecYEG complex, and (ii) dividing the number of micelles by the number of the proteoliposomes in the confocal volume we arrived at the number of SecYEG copies per proteoliposome.

### High-Speed Atomic Force Microscopy (HS-AFM)

HS-AFM imaging (RIBM, Japan) was performed in measuring buffer (50 mM Tris, 50 mM KCl, 50 mM NaCl, 5 mM MgCl2, pH 7.9) at room temperature using USC-F1.2-k0.15 cantilevers (Nanoworld AG, Neuchatel, Switzerland). HS-AFM was operated in tapping mode with free amplitudes of 1.5 – 2.5 nm. The amplitude setpoint was larger than 90 % of the free oscillation amplitude. SecA was either incubated or injected on SecYEG containing periodically suspended membranes (PSMs) which were prepared as previously described.^34^ For incubation, 2 μM SecA was added and rinsed with measuring buffer after 2 min. In the case of injection, SecA was added during HS-AFM imaging to yield final concentrations of 0.4 – 2 μM. The interaction of SecA and SecYEG (change in height values) was analyzed using an in-house tool implemented in MATLAB (Mathworks, Natick, MA). To compile the histograms depicted in Fig. 2, we analyzed 5 independent experiments that included 11 z-traces (such as Fig. 2B) with lengths of up to 85 seconds. We analyzed 151 off- and 157-on events to obtain the respective time constants (Fig. 2C and D).

### Surface Plasmon Resonance

SPR experiments were performed at room temperature on a Horiba OpenPlex SPRi device (Horiba France SAS, Longjumeau, France) and lipophilic LP sensor chips (XANTEC, Duesseldorf, Germany). As running buffer, we used the same measuring buffer as for HS-AFM imaging (50 mM Tris, 50 mM KCl, 50 mM NaCl, 5 mM MgCl2, pH 7.9), but with varying KCl concentrations (10, 50, 200 mM). The flow rate was adjusted to 50 μl/min. Small unilamellar vesicles, prepared from E. coli polar lipid extract (Avanti Polar Lipids, Alabaster, AL) via sonication, were injected at a concentration of 0.2 mg/ml in the respective running buffer for 20 min. After formation of a lipid bilayer on the sensor chip, 500 nM SecA His-ΔN20 was injected, followed by sequential injections of SecA with increasing concentrations (50, 100, 200, 500, 1000 nM). Each injection lasted for 10 min, followed by a 30 min dissociation phase with running buffer. The sensorgrams of these kinetic titration experiments were processed using MATLAB (Mathworks, Natick, MA) and a two state kinetic model as sketched in Fig. 1D was fit to the data.

**Fig. 1.**
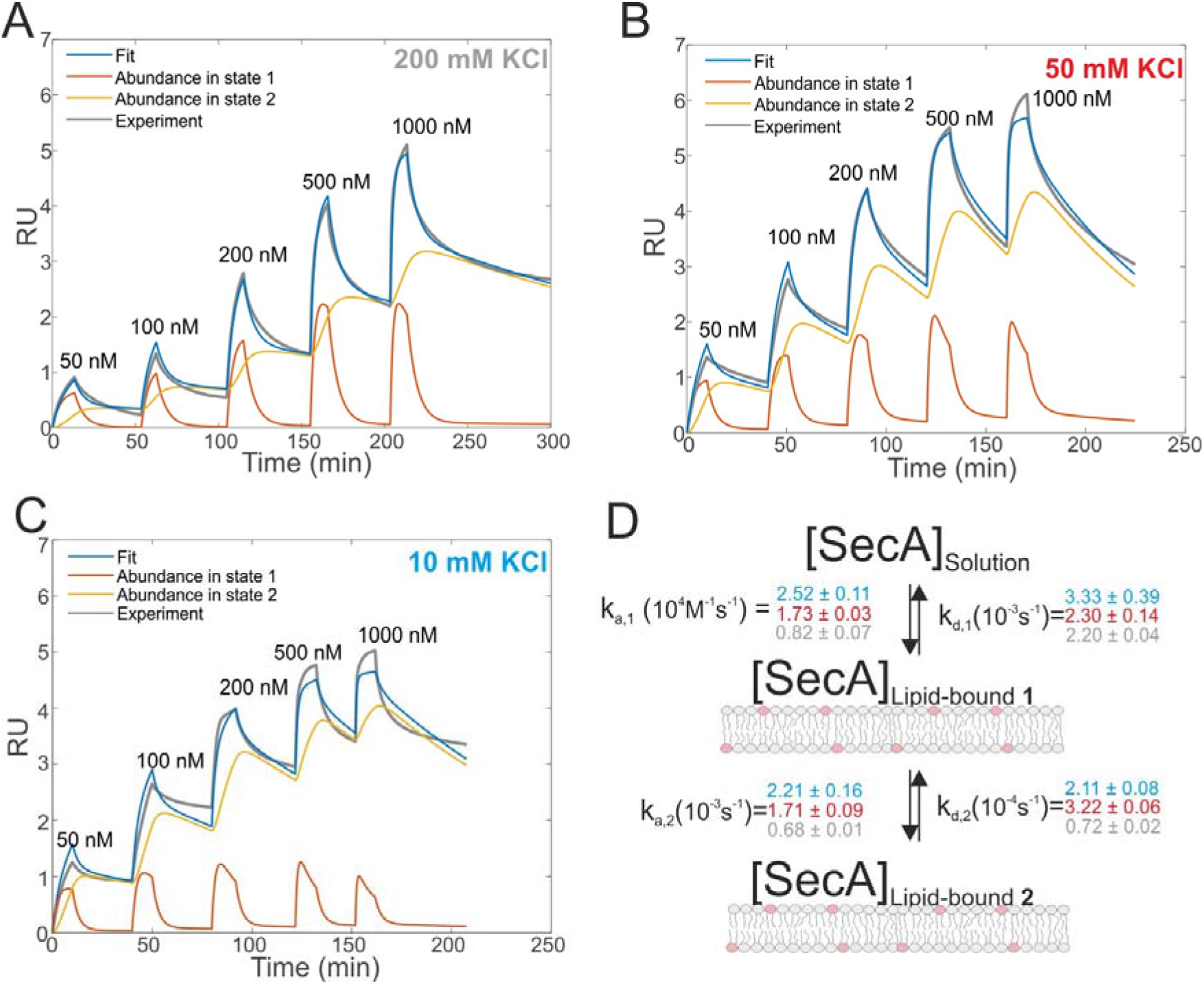
Interaction of SecA with acidic phospholipids containing membranes. **(A)** Surface plasmon resonance experiments of SecA binding to a supported lipid bilayer made of E. coli lipids examined at a KCl concentration of 200 mM. Fit (blue line) represents the sum of abundances in state 1 and 2. **(B)** As in (A) but at 50 mM KCl. **(C)** As in (A) but at 10 mM KCl. **(D)** Sketch of the two state model that best fit the SPR sensorgrams.

## Results

### SecA binds to lipid membranes in a dynamic two-state mechanism

We first studied and quantified the N-terminus mediated interaction of SecA with a lipid membrane in surface plasmon resonance (SPR) experiments. We used the C-terminally truncated SecA mutant N95^35^ that has been used to obtain the structure of the SecA-SecYEG complex.^8^ Small unilamellar vesicles (SUVs) made of *E.coli* lipids were applied to the SPR sensor chip to generate a supported lipid bilayer (SLB). To check for defects in the resulting SLB that potentially cause unspecific adhesion of SecA to the bare dextran surface of the SPR sensor chip, we always applied 500 nM of the SecA His-ΔN20 mutant^7^ prior to the actual experiment and made sure that no binding was observed. We then injected SecA into the SPR flow cell at increasing concentrations (50 – 1000 nM) leading to SecA association, alternated by the injection of pure buffer to induce dissociation, and repeated the experiment at three different KCl concentrations (10, 50, 200 mM) in the running buffer (Fig 1A-C). A two state interaction model best described the resulting SPR sensorgrams: [SecA] [lipids] ⟺ [SecA]_Lipid-bound 1_ ⟺ [SecA lipid]_Lipid-bound 2_ (Fig 1D). The model assumes a monomeric SecA conformation on the membrane surface because (i) complete dissociation of aqueous SecA dimers has previously been observed in the presence of liposomes containing acidic phospholipids^25^, and (ii) artificially stabilized SecA dimers are unable to bind to the membrane surface.^36,37^ The overall SecA abundance on the membrane was comparable at different [KCl], as indicated by the size of the SPR signal (Fig S2). Yet, both the kinetic rates and the relative abundance of SecA in states 1 and 2 that we obtained by fitting the mathematical model to the data strongly varied. At the highest KCl concentrations (200 mM; Fig 1A), the relative abundance of states 1 and 2 was roughly the same. At 50 mM KCl (Fig 1B), state 1 was ~ half as abundant as state 2, and at the lowest KCl concentration (10 mM; Fig 1C) and at the highest SecA concentration (1 μM) state 1 was only 1/4 as abundant as state 2. Apparently, the decreasing salt concentration shifted the equilibrium towards state 2. SecA His-ΔN20 binding to the lipid membrane was no longer negligible (Fig S3) at 10 mM KCl, indicating that the sensorgrams contained additional, N-terminus independent contributions. These unaccounted interactions may explain deviations of the mathematical fit from the sensorgram (Fig 1C).

The kinetic rates obtained from the global fits showed an unique trend (Fig 1D): The initial membrane association rate *k*_*a,1*_ was smallest (0.8 ± 0.1 × 10^4^ M^−1^s^−1^) at the highest salt concentration and increased monotonically with decreasing KCl concentrations (50 mM KCl: 1.7 ± 0.1 × 10^4^ M^−1^s^−1^; 10 mM KCl: 2.5 ± 0.1 × 10^4^ M^−1^s^−1^), indicating that SecA targeting to the membrane (state 1) is driven by electrostatics.

### Direct visualization of two distinct binding modes of SecA to SecYEG

We then performed high speed atomic force microscopy (HS-AFM) experiments to monitor the next step in the molecular pathway, i.e. the encounter and binding of lipid bound SecA to reconstituted SecYEG. We employed periodically suspended membranes (PSMs), which are flat lipid membranes made of *E. coli* liposomes supported by a two-dimensional streptavidin crystal. The latter allows for tuning the lateral mobility of embedded membrane proteins while largely suppressing unwanted interactions with the support.^34^ PSMs are particularly suited for HS-AFM investigations, because the valleys between streptavidin hills may accommodate SecYEG’s periplasmic loops. Being unable to freely migrate between the valleys, SecYEG can easily be imaged by HS-AFM (Movie S1). After inspection of the PSMs and identification of individual SecYEG molecules, which typically protrude up to 2 nm from the membrane, SecA was injected into the HS-AFM liquid cell. After an initial lag time, fast diffusing membrane bound SecA molecules (visible as ‘spike like noise’)^34,38^ and binding events of individual SecA molecules to properly oriented SecYEG complexes (i.e. with their cytoplasmic side facing upwards) were observed (Fig 2A, Movie S2; Additional movies from independent experiments: Movies S3-S4).

**Fig. 2.**
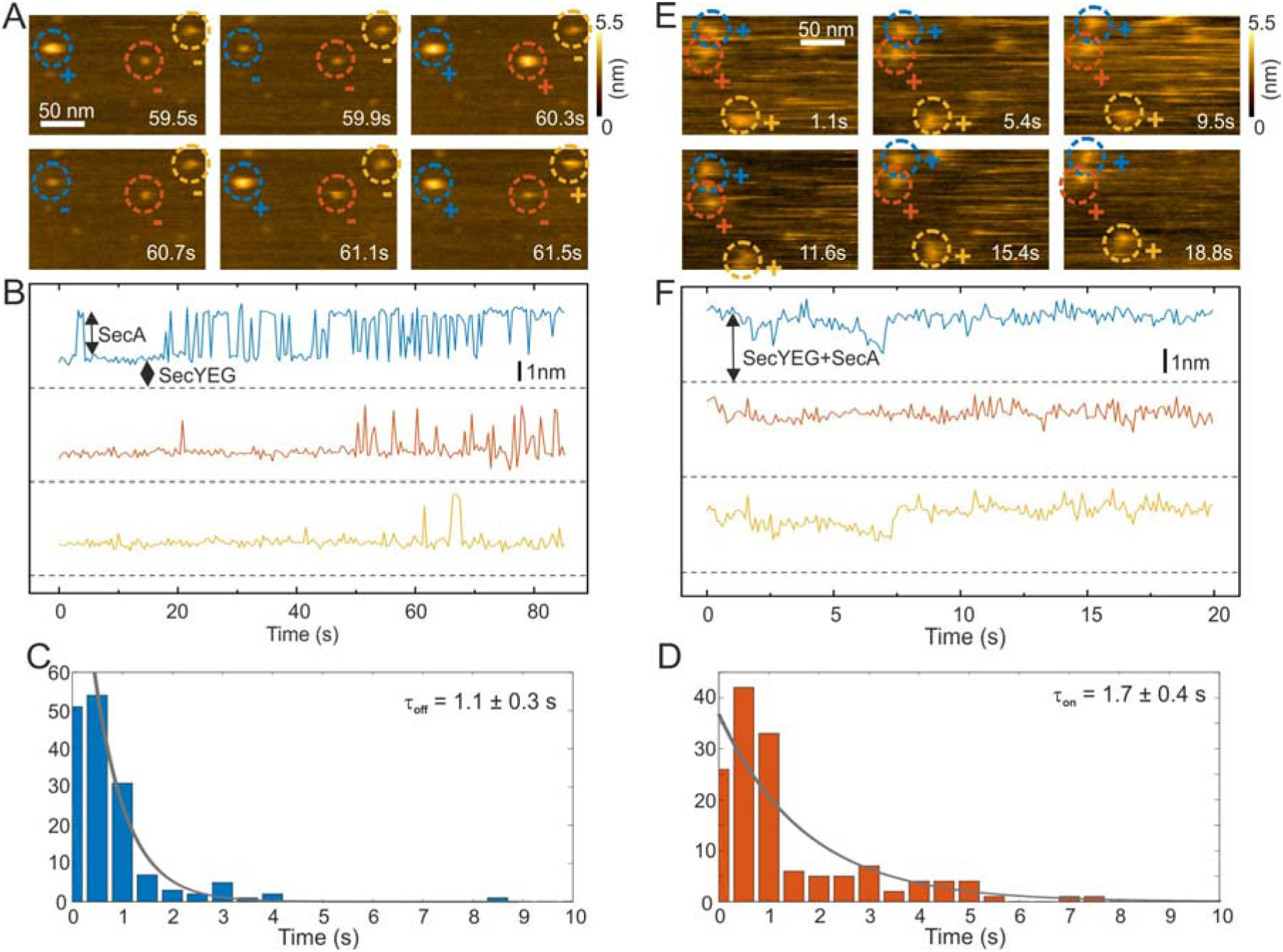
HS-AFM imaging and statistical analysis of SecA binding to SecYEG embedded in partially suspended membranes (PSMs). **(A)** HS-AFM time series (Movie S2) of SecA binding to SecYEG embedded in PSMs. At low SecA concentrations on PSMs, SecA binding to SecYEG was not stationary but exhibited characteristic on/off kinetics. **(B)** Height vs. time traces generated from (A). **(C)** Statistical analysis of SecA-resident times on SecYEG contained in the height vs. time traces (B) and corresponding exponential fit. **(D)** Same as (C), but assessment of timespans until SecA rebinding occurs. **(E)** HS-AFM time series (Movie S5) taken at elevated SecA surface concentration. SecA binding to SecYEG was stationary lacking the kinetics as observed in (A) and (B). **(F)** Height vs. time traces generated from (E).

The SecYEG-SecA complex exhibited a height of up to 5 nm with respect to the surrounding bilayer in accordance with previous (HS)-AFM observations of SecYEG-bound SecA monomers.^34,39^ Remarkably, individual SecA molecules were not stationary bound to the underlying SecYEG but exhibited a discrete on-and off-hopping. Fig 2B exemplifies this dynamic interaction in form of height vs. time traces that were generated from Movie S2. The movie shows the height fluctuation at the position of a SecYEG channel due to the presence/absence of a SecA molecule on top. The traces also highlight that SecA binding was not randomly distributed among the SecYEG channels in the field of view. Rather, the fluctuations localized to the very same SecYEG complex that initially had bound a SecA molecule. The observation suggests trapping of SecA molecules in the vicinity of SecYEG subsequent to SecA dissociation from SecYEG. This behavior was observed in the beginning of the experiments where the membrane bound SecA concentration was low. A statistical analysis of the time periods in which SecA was bound (Fig 2C) or unbound (Fig 2D) yielded characteristic time constants *τ*_off_=1.1 ± 0.3 s and *τ*_on_=1.7 ± 0.4 s, respectively. During the course of the experiment, more and more SecA molecules bound to the membrane and became visible as more pronounced (as compared to the initial phase) topographical (‘spike-like’) fluctuations, representative for fast moving membrane bound objects^34,38^ that were clearly not present when no SecA was added to the HS-AFM liquid cell (cf. Movie S1). At these higher surface concentrations of SecA (Fig 2E, Movie S5) the SecYEG-SecA complexes were stationary, i.e. the initial on-and off-hoping was not observed anymore, which is also evident from the corresponding height vs. time traces (Fig 2F). Taken together, the HS-AFM experiments suggest three different states of SecA associated with SecYEG: (I) A state where SecA is localized in the vicinity of SecYEG but not directly bound to it (effective lifetime *τ*_on_ after initial encounter, may be associated with a lipid-annulus surrounding SecYEG^40^). SecA may be too loosely attached to be imaged via HS-AFM, (II) A low affinity state where SecA is loosely bound to SecYEG (lifetime *τ*_off_), and (III) A high affinity state where SecA forms a stable complex with SecYEG (Fig 2E, F). Notably, the HS-AFM movies did not provide evidence for distinct structural differences between states II and III, as the differences in height or lateral dimensions of the respective SecA-SecYEG complexes were not resolvable (Fig 2A vs. E).

### Structural characterization of SecA binding to SecYEG

To further characterize and structurally distinguish the low and high-affinity SecYEG-bound states observed in HS-AFM experiments, we performed luminescence resonance energy transfer (LRET) experiments between a fluorescent dye (ATTO488) on SecA’s two helix finger (K797; acceptor) and a terbium ion (donor) in a genetically engineered lanthanide binding pocket (LBT)^15,27^ inserted at position G297 within helix-connecting loops at the periplasmic face of SecYEG, respectively (Fig 3A, Fig. S4). Using terbium as donor molecule limits the maximal positional error to −11% +12% due to the isotropic emission of terbium ions^29^ compared to +− 100% in case of FRET experiments with a second organic dye. The positions on SecA and SecYEG were chosen to accurately measure the insertion depth of the two-helix finger into the SecYEG channel when SecA is engaged with the translocon.

**Fig. 3.**
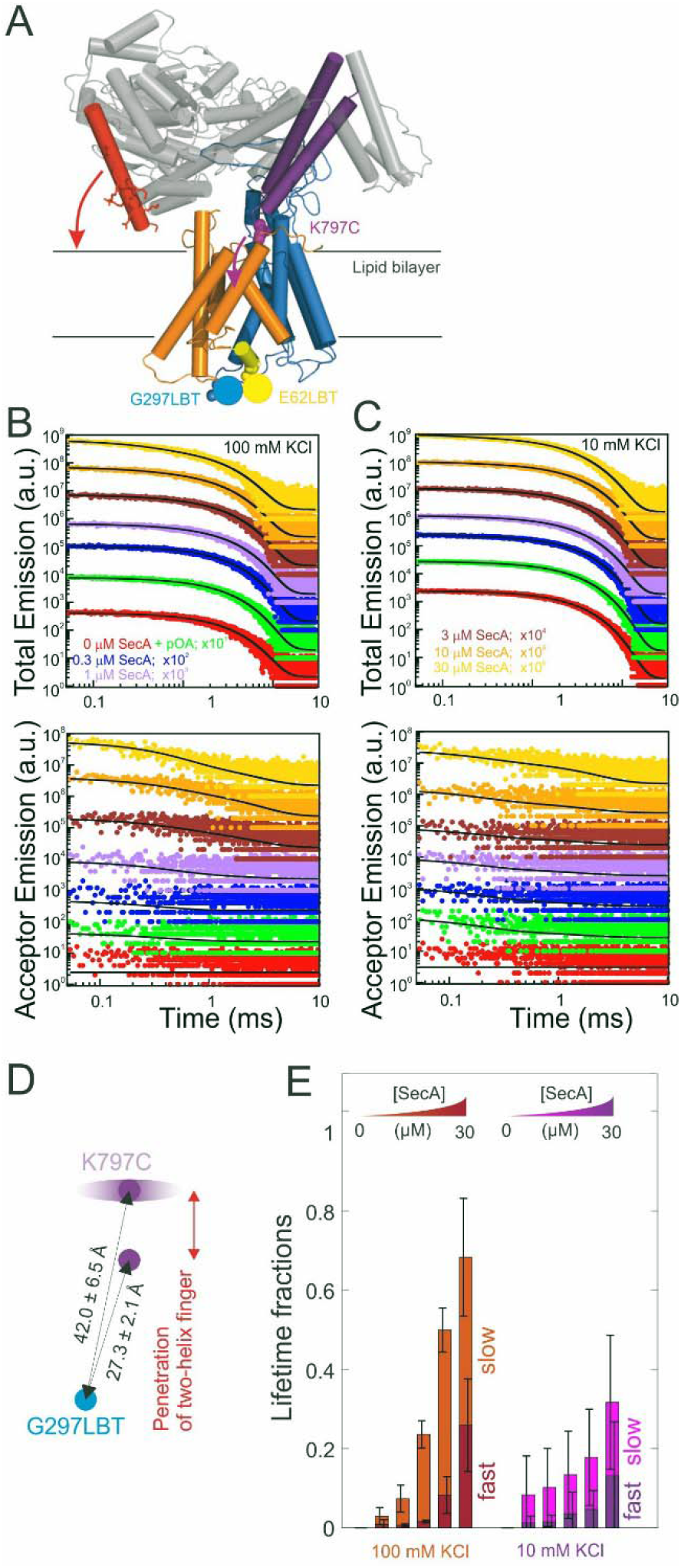
LRET structural assessment of distinct SecA-SecYEG complexes. **(A)** Localization of the LBT tags and the fluorescent dyes in the SecY-SecA complex. SecY (orange and blue; emanating from the lateral gate) residues E62 (plug) and G297 (periplasmic loop) are marked in yellow and blue, respectively. SecA’s two-helix finger is displayed in purple and its N-terminus is colored in red. Arrows indicate the potential domain movements. **(B)** Total fluorescence decay curves (upper panel, double logarithmic plot) and acceptor emission curves (lower panel, double logarithmic plot) of reconstituted SecYEG G297LBT incubated with increasing amounts of ATTO488 labeled SecA recorded at 100 mM KCl. As denoted in the legend, curves were vertically shifted (multiplied by 10^n^) for better visualization. Black curves indicate the result of the global fitting procedure which yielded fluorescence lifetimes *τ*_*D*_ = 2386 μs, *τ*_*T, fast*_ = 337 μs, and *τ*_*T, slow*_ = 4954 μs. **(C)** Same as in (B) but recorded at 10 mM KCl. Black curves indicate the result of the global fitting procedure which yielded fluorescence lifetimes TD = 2472 μs, *τ*_*T, fast*_ = 145 μs, and *τ*_*T, slow*_ = 2876 μs. **(D)** Distances between the G297LBT and the tip of the two-helix finger calculated from the lifetimes in (E). Two distinct structural states of the complex were observed, differing in the degree of penetration of SecA’s two-helix finger into the translocation channel. **(E)** Fractions of lifetimes from (B) and (C) (Averages from n=4 independent experiments). Addition of proOmpA prior to SecA didn’t show any effect, binding of SecA to SecYEG was enhanced at physiological salt conditions (100 mM). Two distinct SecA-SecYEG complexes differing in the penetration depth of SecA’s two-helix finger into the SecY channel were registered in a detailed analysis of fluorescence decay curves (C) and (D). The errors arise from averaging of multiple experiments. Obtained distances for 100mM and 10 mM titrations were pooled (n=8).

In detergent, SecY G297LBT showed only a very weak, salt-independent interaction with SecA K797C-ATTO488, as evident from the small changes in total emission and acceptor emission decay curves upon SecA addition (Fig S5A and B). The occupancy of SecYEG molecules by nucleotide-free SecA amounted to about 10 %. This is in agreement with the reported requirement for ATP to form stable complexes in detergent.^8^ The relaxation of donor fluorescence intensity was best described by a model taking into account a single acceptor lifetime. The latter corresponded to a distance of 28.5 ± 0.1 Å between the G297LBT and the tip of the two helix finger. This distance is much shorter than that displayed in the crystal structure of the SecYEG - SecA complex, i.e. it indicates a deep penetration of the two helix finger into the translocation channel. This observation suggests that nucleotide release (or hydrolysis) may trigger the movement of the two-helix finger into the SecY pore. Such movement would be compatible with the previously published power stroke mechanism, i.e. with the two-helix finger pushing the polypeptide chain.^7,8^

Reconstitution of SecYEG G297LBT into proteoliposomes resulted in a much more robust binding of labeled SecA. The decay time of fluorescence intensity depended on both the SecA and KCl concentrations (Fig 3B and C, 100 and 10 mM KCl, respectively; total emitted fluorescence, upper panel; acceptor fluorescence, lower panel). In contrast to the measurements performed in detergent, these data were globally best fit by a model taking into account two distinct acceptor fluorescence lifetimes (cf. materials and methods). Consequently, the distances between the G297LBT and the tip of the two-helix finger calculated from these lifetimes, 27.3 ± 2.1 Å and 42.0 ± 6.5 Å represent two distinct structural states of the complex, differing in the degree of penetration of SecA’s two-helix finger into the translocation channel (Fig 3D). Apparently, the smaller distance matches the distance and thus the conformation determined in detergent, but the larger distance only appears in the presence of a lipid bilayer or when nucleotides are bound.^8^

SecYEG occupancy increased monotonically with increasing SecA concentration. This change was accompanied by an increase in the SecA fraction that elicited the fast relaxation of fluorescence due to the deep penetration of the two-helix finger. At maximum SecA concentrations, the sizes of both slow and fast relaxing fractions were almost equal to each other (Fig 3E). Replication of these experiments with the LBT located at an alternative position within SecY (E62 on the so-called ‘plug’), essentially confirmed the existence of two SecY-insertion distinct states of the two-helix finger in the SecA-SecYEG complex (Fig S6).

SecA-SecYEG interaction was salt dependent. At low KCl concentrations, the fraction with a deep penetration of the two-helix finger was small (~ 20%). It monotonically increased with increasing KCl concentrations (Fig S5C-F recorded at saturating SecA concentrations; Fig 5E, 100 mM vs. 10 mM at different SecA concentrations). That is, the fraction of donor molecules that exhibit a fast relaxation to the ground state increased with KCl concentration. It became equal to the fraction of donors with slow relaxation times at 100 mM KCl. We conclude that the reduction of the effective membrane surface potential at higher ionic strength shifts the equilibrium towards the SecA conformation of with a deeply inserted two-helix finger.

### Ion channel activity of the SecA - SecYEG complex

To check whether the distinct states of bound SecA physically open the translocon, we recorded SecYEG’s single ion channel activity, i.e. performed electrophysiological experiments. We added SecYEG-proteoliposomes to the hypertonic solution (500 mM KCl) at one side of a pre-formed planar lipid bilayer. The solution at the other side was hypotonic (100 mM KCl). The resulting transmembrane osmotic gradient served to fuse vesicles with open SecYEG complexes to the bilayer.^18^ In agreement with previous data,^41^ no channel activity was observed in the absence of SecA (Fig. S7). Two types of channel activity were observed in the presence of SecA: short living channels with a comparatively small unitary conductance (Fig 4A) and long living fully opened SecYEG complexes with a larger conductance (Fig 4B). To illustrate the large difference between both types of channels, we provide (i) a record where a small channel is opening on top of a larger one that most of the time stays open (Fig 4C and D) and a bar chart that visualizes the differences in amplitude (Fig 4C,inset). The smaller events are likely to represent channels that are primed to receive the polypeptide chain but have not yet fully opened, i.e. they may correspond to those channels that were captured by the crystal structure of the SecYEG complex with SecA.^8^ In contrast, the events of large conductivity *G* of 600±80 pS are similar in size to the ion channels observed after plug deletion^17^ or ribosome binding.^19^

**Fig. 4.**
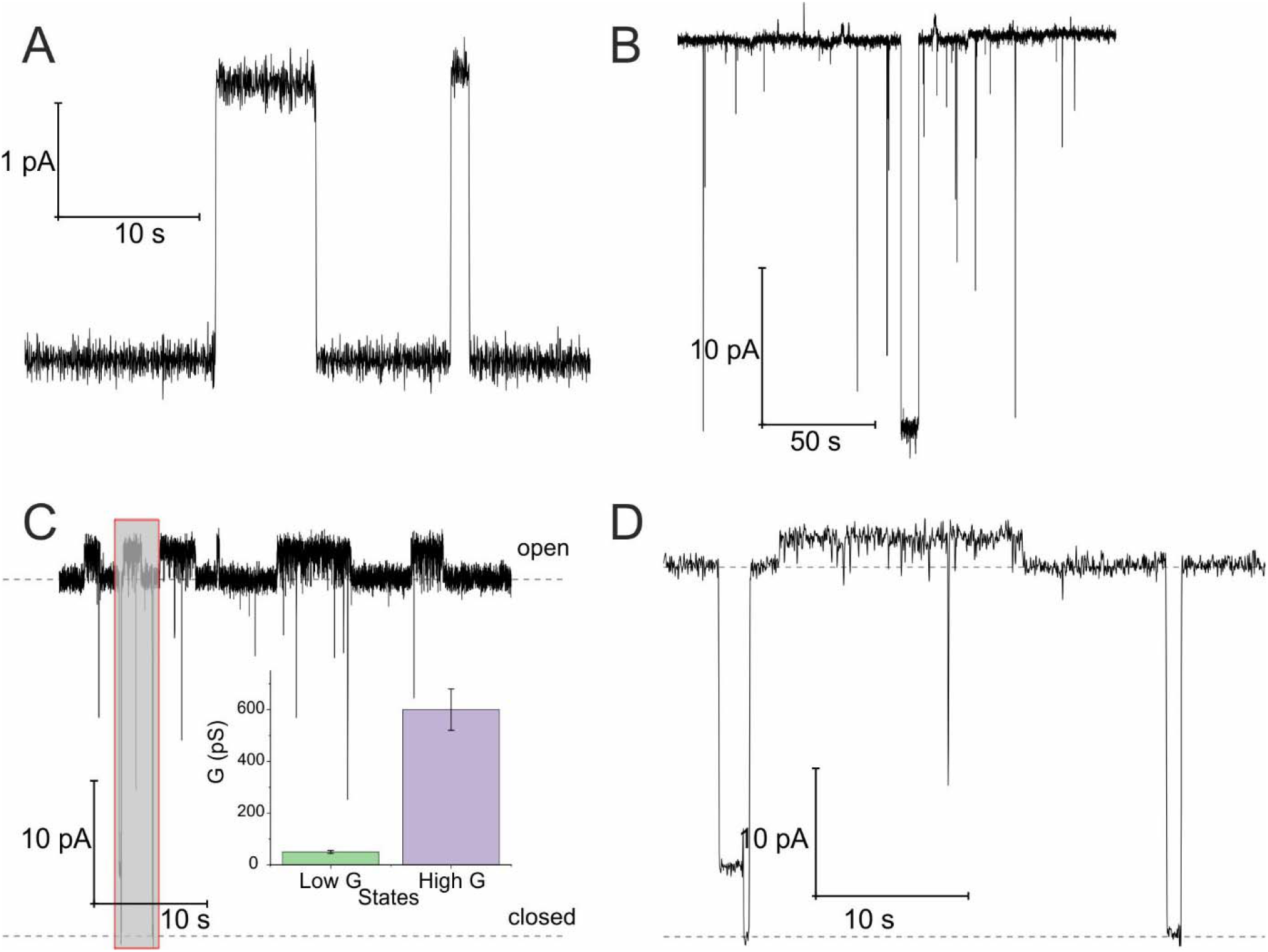
Opening of the SecYEG translocon upon SecA binding. Two distinct conductivity states of the SecYEG-SecA complex are observed. **(A)** Short living low conductivity states are characterized by an amplitude of 2.5 pA. **(B)** A longer living high conductivity state with an amplitude of ~ 30 pA. **(C)** Record of the simultaneous activity of both channel types. It illustrates amplitude and lifetime differences between the large predominantly open channels and the smaller channels with a lifetime of a few seconds. Inset: Bar chart of conductivities. The concentration of SecA in the hypertonic compartment was 2.1 μM. The transmembrane potential amounted to 50 mV. **(D)** Display of the grey fragment of the record in (C) at an extended time scale. The lower dashed line corresponds to the level of background current where the channels are closed. The upper dashed line visualizes the level of current through a single fully open channel.

## Discussion

The nucleotide-free SecA monomer interacts with both the SecYEG complex and lipid membranes. When taken separately, binding to both partners is weak. Yet, when acting in unison, the lifetime *τ* of the SecA - SecYEG complex may reach tens of seconds. Assuming a diffusion limited on-rate *k*_on_, the observation suggests an apparent equilibrium binding constant *K*_*app*_ of the nucleotide-free SecA to the reconstituted SecYEG complex in the nanomolar range:

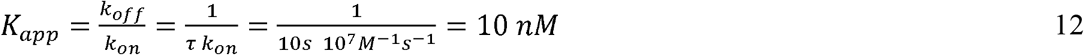

where the release constant *k*_off_ is equal to 1/*τ*. Below we perform a quantitative analysis of the two separate binding events in order to obtain mechanistic insight into the molecular origin of the astonishing K value.

Binding to membranes containing acidic phospholipids emerges as a dynamic two-state process. It involves the transition between an initially formed complex (Fig 5, State 1) to a more tightly bound complex (State 2) as demonstrated by SPR experiments. The kinetic on-rate (k_a,1_) of this interaction decreases with increasing salt concentrations suggesting that the initial driving force of SecA membrane targeting and therefore its interaction with the membrane in state 1 is of electrostatic nature.^3^ The transition between state I and II may be associated with the insertion of the N-terminus into the lipid membrane as previously suggested.^4^

**Fig. 5.**
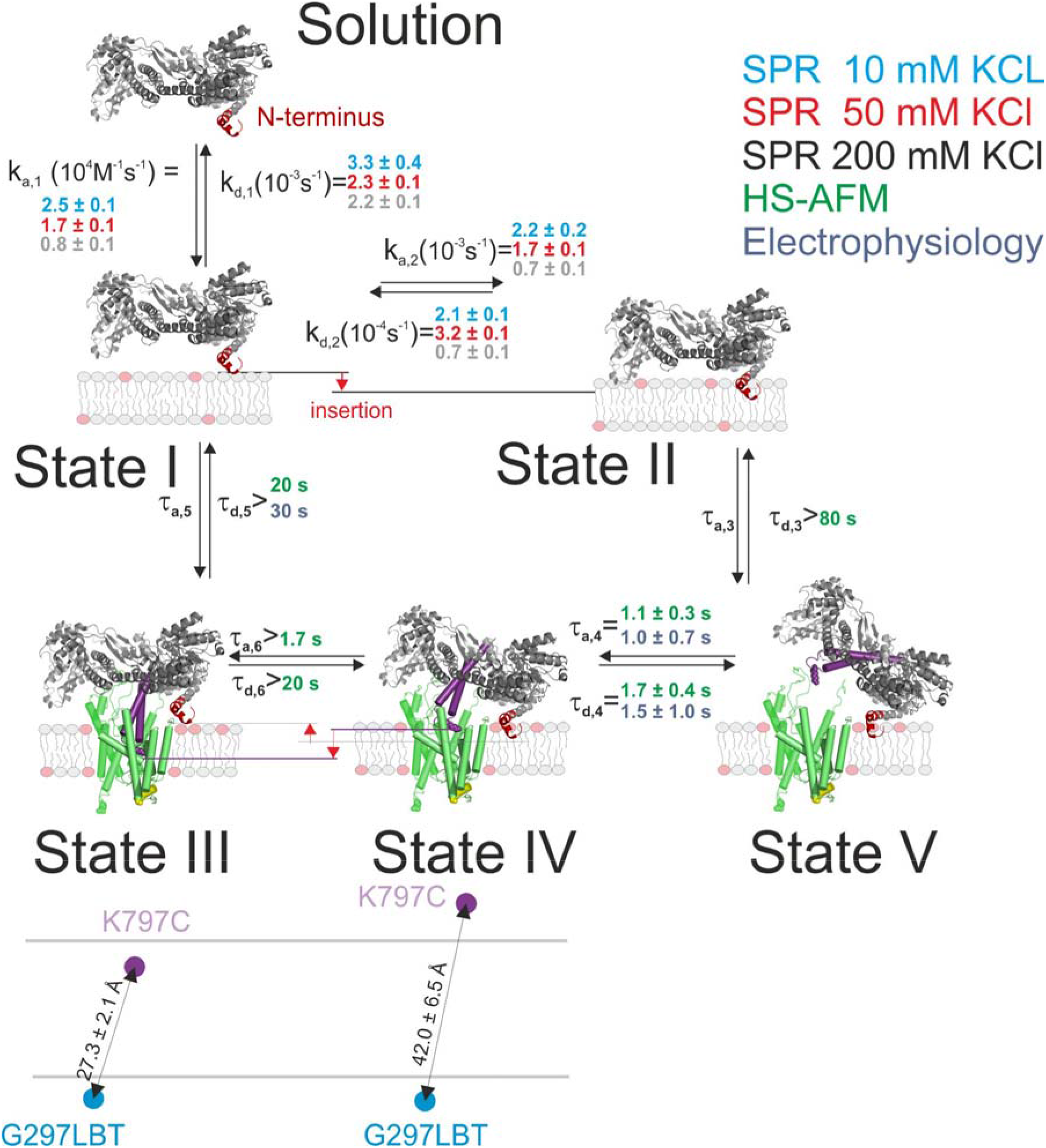
Kinetic model of SecA targeting SecYEG via the lipid membrane. Kinetic rates and lower boundaries of life-times were taken from the experiments depicted in Figs. 1–4.

The model does not refer to the SecA dimer, because it is (i) apparently unable to bind to the membrane surface^36,37^, and (ii) dissociates the presence of acidic lipids^25^. Reports about the functionality of covalently linked SecA dimers^5,42^ do not challenge the conclusion, because they do not contain evidence that the dimer may bind to the lipid in the absence of SecYEG. Truncation of residues at the N-terminus may hamper SecA dimerization.^5^ Yet, the conclusion that the dimer is essential for protein translocation is not justified.^6^ Rather, the deletion affects the binding of the SecA monomer to the lipid membrane ^7^, thereby grossly minimizing the probability of SecA to find the translocation channel. This interpretation is also in line with our HS-AFM images. They show a SecA monomer on top of the SecYEG molecule. The SecA dimensions are clearly incompatible with dimeric structures of the protein.

HS-AFM observations of SecA binding to membrane embedded SecYEG indicate the existence of high and low affinity SecA-SecYEG complexes. The high affinity complex (State III) did not dissociate within the time frame of the experiments, suggesting lifetimes (*τ*_d,5_, *τ*_d,6_) of tens of seconds or even minutes. In contrast, the low affinity complex (State IV) frequently dissociated (*τ*_d,4_). It ended up in a defined state (State V) close to the SecYEG channel. Re-association events (*τ*_a,4_) occurred (Fig 2A and B). One may speculate that in state V SecA is associated with the acidic phospholipid annulus surrounding SecYEG by inserting its N-terminus into the membrane. Interestingly, the hopping between states IV and V did neither lead to a transition into state III (stable binding) nor into the formation of a sole lipid bound state (such as state II). Accordingly, only lower limits for the lifetime in state IV (*τ*_a,6_, *τ*_d,3_) were derived from the respective observation times.

LRET experiments showed the existence of one (in detergent; in the absence of a lipid bilayer) or two (when SecYEG is reconstituted into a lipid bilayer) structurally distinct SecA-SecYEG complexes (Fig 5; States III and IV). They differ in the penetration depth of the two-helix finger. The observation is in line with single molecule FRET experiments in which the movement of the two helix finger was interpreted in terms of a power stroke.^43^ The model envisions the finger to push the polypeptide into the channel. Its retrograde movement does not lead to backsliding, because the clamp domain of SecA tightens around the polypeptide during ATP hydrolysis. Importantly, the first of the two penetration depths that LRET revealed for SecA’s two-helix finger coincided for detergent-solved SecYEG and membrane-embedded SecYEG complexes (State III). However the second penetration depth was not observed in detergent, further supporting the idea that State IV is strongly associated with the surrounding lipid annulus and a potential insertion of SecA’s N-terminus therein.

Single ion channel recordings also supported the existence of two states: two classes of channels with distinct unitary ion conductivities were observed. Their open times were roughly similar to the lifetimes of the SecA – SecYEG complexes as observed in HS-AFM experiments, thereby assigning the observed conductivity states to the respective structural states. It is important to note that the observation of these channels is only possible in partly de-energized membranes. That is, physiological values of the membrane potential serve to close the ion channels^18^, thereby restoring the membrane barrier to ions and small molecules.^44^

Our model suggests that the insertion of SecA’s N-terminus into the lipid membrane hinders a deep penetration of SecA’s two-helix finger into SecYEG. Thus, SecA is trapped shuttling in between state IV and V. However, it does not require a major conformational change^2,4^ to strongly bind to SecYEG as in state III the N-terminus is functioning solely as membrane-tether^7^ that is electrostatically linked to the membrane, but not inserted into it.

The strong binding must be the result of (i) hydrophobic interactions between the two helix finger with the SecYEG channel (equilibrium dissociation constant K_SecYEG_) and (ii) electrostatic interactions between the N-terminus and the membrane (as in lipid bound state I; Equilibrium dissociation constant 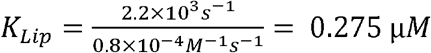. The combined equilibrium dissociation constant of such a hetero-bivalent interaction reads^45,46^:

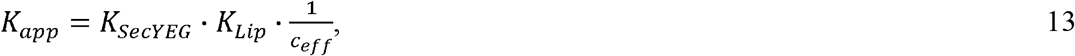

with *c*_*eff*_ being the effective concentration for rebinding when one of the two interactions dissociates. We can use this equation together with 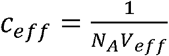 (with *V*_*eff*_ being modelled as a half sphere with a radius of 10 nm) and *K*_*app*_ to estimate *K*_*SecYEG*_ = 30 μM. The SecA-SecYEG binding affinity in the absence of lipids and ATP is thus 8 times lower than the binding affinity of detergent-solubilized SecYEG in the presence of ATP (3.9 μM),^47^ which is likely the reason why the crystallization of the ATP-free SecA-SecYEG complex was not achieved.^8^

If SecA’s affinity to the membrane embedded SecYEG would be equally weak (=*K*_SecYEG_), the motor molecule would be released subsequent to every ATP hydrolysis cycle. The presence of a translocating polypeptide would not preserve the SecYEG-SecA complex, because phosphate release opens SecA’s clamp so that the polypeptide chain passively slides through SecYEG and through SecA.^8,43^ In the interval between the release of one SecA molecule and the binding of a new SecA molecule, the translocating polypeptide would slide back, thereby significantly reducing the translocation efficiency. In contrast, the now observed *K*_app_ = 10 nM ensures a SecA residence time that easily bridges the comparatively short time window between ADP release and ATP capture. As a result, one SecA may perform ~ 260 ATP cycles^7^ before being released. Assuming a hydrolysis rate of 7.6 s^−1^ ^47^ we estimate an average residence time of ~34 s. The overestimated frequency of SecA-exchanges reported by single molecule experiments^43^ may be due to blinking events of the involved fluorophores.^48^

## Conclusion

The interaction between the membrane embedded translocation channel SecYEG and the nucleotide-free SecA monomer is a representation of a unifying principle known from many other high-affinity interactions: the amplification of binding affinity by dual attraction of two interaction sites on a single ligand to two neighboring sites of the binding partner. As a result, the SecYEG - SecA complex is stable enough to survive the interval between ADP release and ATP binding, enabling SecA to perform multiple cycles of hydrolysis while bound to SecYEG.

## Author Contributions

J.P. and P.P. conceived the project. K.W. and A.H. established the LRET setup, performed LRET experiments. K.W., A.H., and C. H. analyzed data. A.K. performed HS-AFM and SPR experiments and analyzed data. D.K. performed single channel electrophysiology experiments and analyzed data. R.K., C.S., M.Z., and K.W. expressed, purified, reconstituted proteins and performed translocation assays. J.P., A.H., and P.P. wrote the manuscript. All authors commented on the manuscript.

## Acknowledgments

This work was supported by the Austrian Science Fund (FWF, P25844 to J.P. and P28213 to PP), the European Fund for Regional Development (EFRE, IWB2020) and the Federal State of Upper Austria.

