## Supplementary Figures for "Interaction of the Motor Protein SecA and the Bacterial Protein Translocation Channel SecYEG in the Absence of ATP"

Electronic Supplementary Information

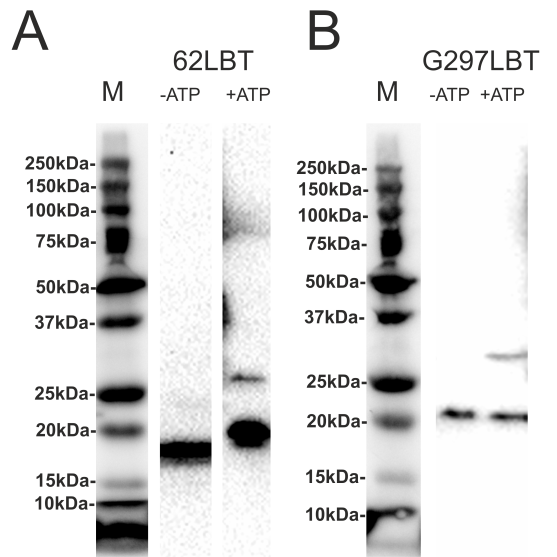

**Figure S1. Translocation Assay.** Anti-His-HRP (Milenitec) detection shows translocated pOD at 27 kDA in ATP positive samples for SecYEG with mutations 62LBT (A) or 297LBT (B).

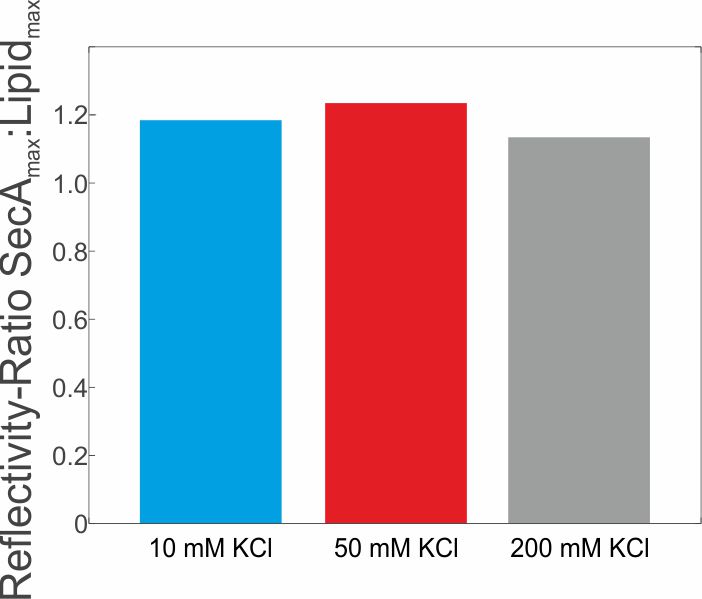

**Figure S2. Maximal SecA to lipid mass ratio on the SPR sensor chips.** Ratio of resonance units of total lipids adsorbed to the SPR chip and the maximal amount of lipid bound SecA (at 1000 nM concentration).

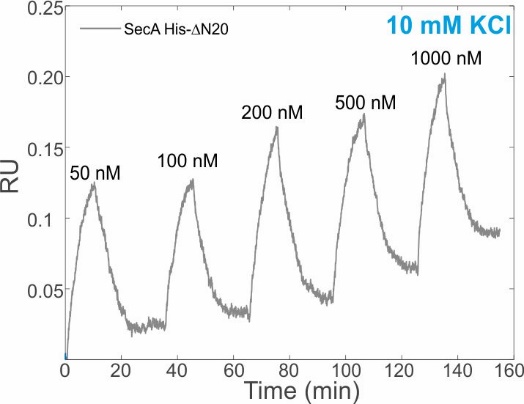

**Figure S3. Non-N-terminus mediated interaction of SecA at low salt concentration**. Surface plamson resonance senorgram of a concentration series of His-ΔN20 binding to a supported lipid bilayer deposited on a SPR sensor chip at 10 mM KCl.

| SecA(N95) K797C  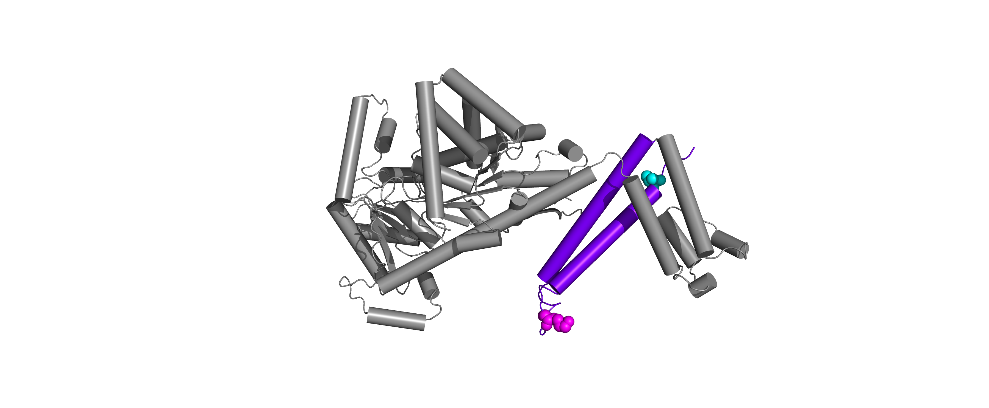 | MLIKLLTKVFGSRNDRTLRRMRKVVNIINAMEPEMEKLSDEELKGKTAEFRARLEKGEVLENLIPEAFAVVREASKRVFGMRHFDVQLLGGMVLNERSIAEMRTGEGKTLTATLPAYLNALTGKGVHVVTVNDYLAQRDAENNRPLFEFLGLTVGINLPGMPAPAKREAYAADITYGTNNEYGFDYLRDNMAFSPEERVQRKLHYALVDEVDSILIDEARTPLIISGPAEDSSEMYKRVNKIIPHLIRQEKEDSETFQGEGHFSVDEKSRQVNLTERGLVLIEELLVKEGIMDEGESLYSPANIMLMHHVTAALRAHALFTRDVDYIVKDGEVIIVDEHTGRTMQGRRWSDGLHQAVEAKEGVQIQNENQTLASITFQNYFRLYEKLAGMTGTADTEAFEFSSIYKLDTVVVPTNRPMIRKDLPDLVYMTEAEKIQAIIEDIKERTAKGQPVLVGTISIEKSELVSNELTKAGIKHNVLNAKFHANEAAIVAQAGYPAAVTIATNMAGRGTDIVLGGSWQAEVAALENPTAEQIEKIKADWQVRHDAVLEAGGLHIIGTERHESRRIDNQLRGRSGRQGDAGSSRFYLSMEDALMRIFASDRVSGMMRKLGMKPGEAIEHPWVTKAIANAQRKVESRNFDIRKQLLEYDDVANDQRRAIYSQRNELLDVSDVSETINSIREDVFKATIDAYIPPQSLEEMWDIPGLQERLKNDFDLDLPIAEWLDKEPELHEETLRERILAQSIEVYQRKEEVVGAEMMRHFEKGVMLQTLDSLWKEHLAAMDYLRQGIHLRGYAQ**C**DPKQEYKRESFSMFAAMLESLKYEVISTLSKVQVHHHHHH |
| --- | --- |
| SecA(N95) K797C V829LBT | MLIKLLTKVFGSRNDRTLRRMRKVVNIINAMEPEMEKLSDEELKGKTAEFRARLEKGEVLENLIPEAFAVVREASKRVFGMRHFDVQLLGGMVLNERSIAEMRTGEGKTLTATLPAYLNALTGKGVHVVTVNDYLAQRDAENNRPLFEFLGLTVGINLPGMPAPAKREAYAADITYGTNNEYGFDYLRDNMAFSPEERVQRKLHYALVDEVDSILIDEARTPLIISGPAEDSSEMYKRVNKIIPHLIRQEKEDSETFQGEGHFSVDEKSRQVNLTERGLVLIEELLVKEGIMDEGESLYSPANIMLMHHVTAALRAHALFTRDVDYIVKDGEVIIVDEHTGRTMQGRRWSDGLHQAVEAKEGVQIQNENQTLASITFQNYFRLYEKLAGMTGTADTEAFEFSSIYKLDTVVVPTNRPMIRKDLPDLVYMTEAEKIQAIIEDIKERTAKGQPVLVGTISIEKSELVSNELTKAGIKHNVLNAKFHANEAAIVAQAGYPAAVTIATNMAGRGTDIVLGGSWQAEVAALENPTAEQIEKIKADWQVRHDAVLEAGGLHIIGTERHESRRIDNQLRGRSGRQGDAGSSRFYLSMEDALMRIFASDRVSGMMRKLGMKPGEAIEHPWVTKAIANAQRKVESRNFDIRKQLLEYDDVANDQRRAIYSQRNELLDVSDVSETINSIREDVFKATIDAYIPPQSLEEMWDIPGLQERLKNDFDLDLPIAEWLDKEPELHEETLRERILAQSIEVYQRKEEVVGAEMMRHFEKGVMLQTLDSLWKEHLAAMDYLRQGIHLRGYAQ**C**DPKQEYKRESFSMFAAMLESLKYEVISTLSKV**YIDTNNDGWYEGDELLA**QVHHHHHH |
| SecY G297LBT  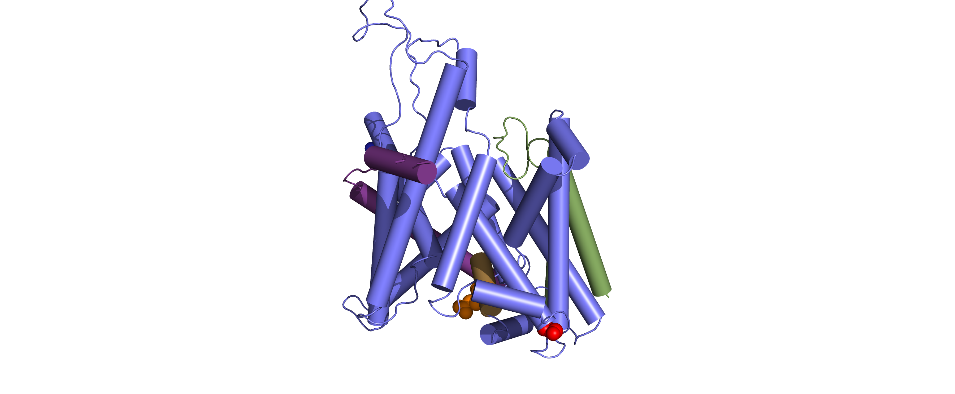 | MAKQPGLDFQSAKGGLGELKRRLLFVIGALIVFRIGSFIPIPGIDAAVLAKLLEQQRGTIIEMFNMFSGGALSRASIFALGIMPYISASIIIQLLTVVHPTLAEIKKEGESGRRKISQYTRYGTLVLAIFQSIGIATGLPNMPGMQGLVINPGFAFYFTAVVSLVTGTMFLMWLGEQITERGIGNGISIIIFAGIVAGLPPAIAHTIEQARQGDLHFLVLLLVAVLVFAVTFFVVFVERGQRRIVVNYAKRQQGRRVYAAQSTHLPLKVNMAGVIPAIFASSIILFPATIASWFGGG**YIDTNNDGWYEGDELLA**TGWNWLTTISLYLQPGQPLYVLLYASAIIFFSFFYTALVFNPRETADNLKKSGAFVPGIRPGEQTAKYIDKVMTRLTLVGALYITFISLIPEFMRDAMKVPFYFGGTSLLIVVVVIMDFMAQVQTLMMSSQYESALKKANLKGYGR |
| SecY E62LBT | MAKQPGLDFQSAKGGLGELKRRLLFVIGALIVFRIGSFIPIPGIDAAVLAKLLEQQRGTIIE**YIDTNNDGWYEGDELLA**MFNMFSGGALSRASIFALGIMPYISASIIIQLLTVVHPTLAEIKKEGESGRRKISQYTRYGTLVLAIFQSIGIATGLPNMPGMQGLVINPGFAFYFTAVVSLVTGTMFLMWLGEQITERGIGNGISIIIFAGIVAGLPPAIAHTIEQARQGDLHFLVLLLVAVLVFAVTFFVVFVERGQRRIVVNYAKRQQGRRVYAAQSTHLPLKVNMAGVIPAIFASSIILFPATIASWFGGGTGWNWLTTISLYLQPGQPLYVLLYASAIIFFSFFYTALVFNPRETADNLKKSGAFVPGIRPGEQTAKYIDKVMTRLTLVGALYITFISLIPEFMRDAMKVPFYFGGTSLLIVVVVIMDFMAQVQTLMMSSQYESALKKANLKGYGR |
| SecE | MHHHHHHGGPSANTEAQGSGRGLEAMKWVVVVALLLVAIVGNYLYRDIMLPLRALAVVILIAAAGGVALLTTKGKATVAFAREARTEVRKVIWPTRQETLHTTLIVAAVTAVMSLILWGLDGILVRLVSFITGLRF |
| SecG | MYEALLVVFLIVAIGLVGLIMLQQGKGADMGASFGAGASATLFGSSGSGNFMTRMTALLATLFFIISLVLGNINSNKTNKGSEWENLSAPAKTEQTQPAAPAKPTSDIPN |
| proOmpA | MKKTAIAIAVALAGFATVAQAAPKDNTWYTGAKLGWSQYHDTGFINNNGPTHENQLGAGAFGGYQVNPYHHHHHHC |

**Figure S4. Structure and sequence of the used constructs.**

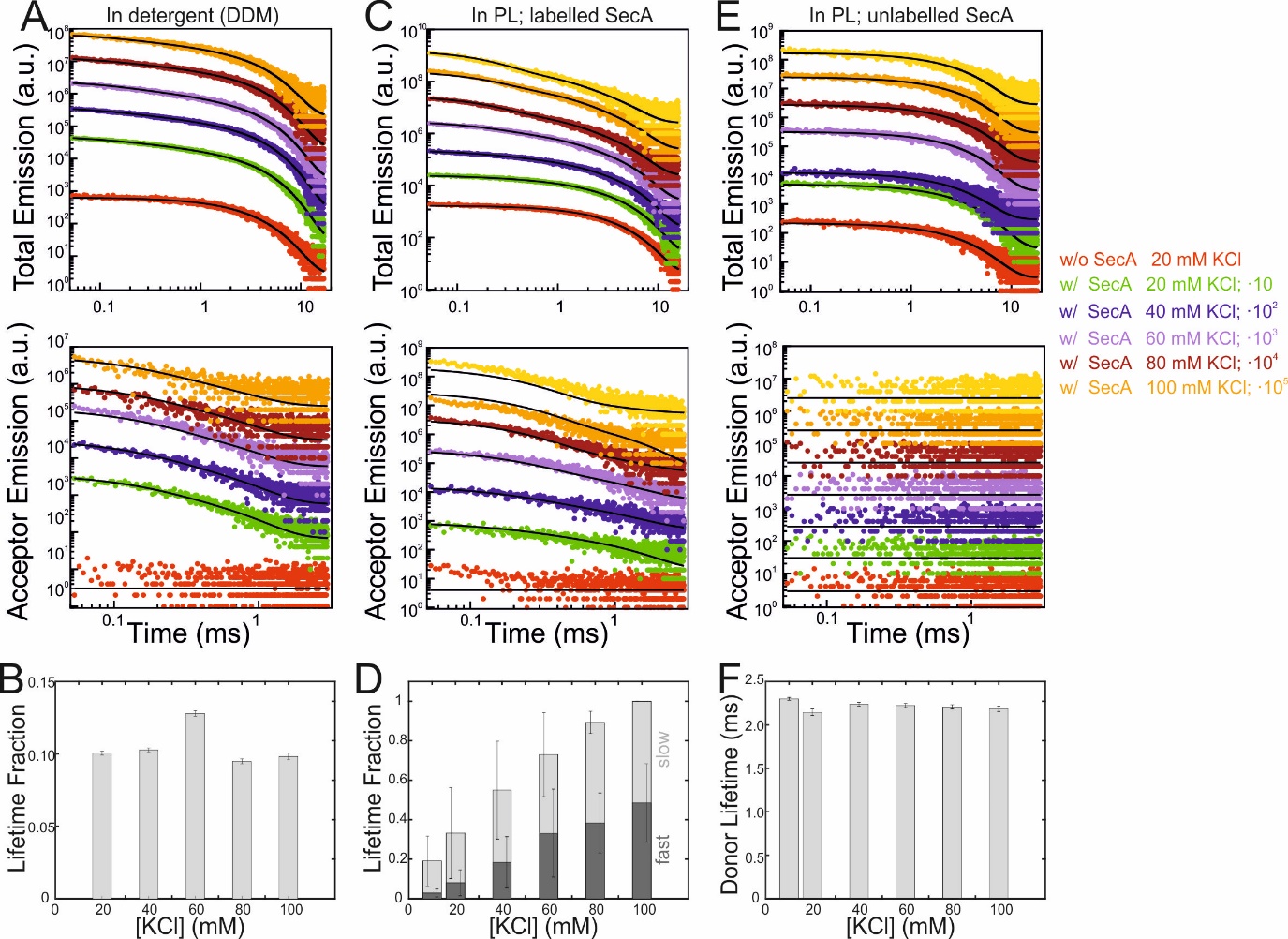

**Figure S5** **Influence of salt on the SecA-SecYEG binding.** **(A)** Total fluorescence decay curves (upper panel) and acceptor emission (lower panel) of 50 µM SecA K797C-Atto488 binding to SecYEG G297LBT in detergent (DDM). As denoted in the legend, curves were vertically shifted (multiplied by 10n) for better visualization. Black curves indicate the result of the global fitting procedure. **(B)** Fraction of acceptor emission from a, obtained from a global single-component acceptor fit. The obtained time constant was 326 ± 3 µs and the resulting donor-acceptor distance 28.5 ± 0.1 Å (error intervals were calculated form the 95% confidence interval of the determined lifetime). **(C)** Same as in (A) but with ATTO532-labeled SecA and SecYEG reconstituted into proteoliposomes. **(D)** Average fractions of fast and slow lifetimes of two experiments as in (C) normalized to 100% at the highest salt concentration (accounting for varying orientations of reconstituted SecYEG). **(E)** Same as in (C) but with unlabeled SecA. Acceptor emission (lower panel) was zero and total emission traces were indistinguishable for all [KCl]. **(F)** Donor lifetimes determined from the traces in (E) suggesting that a salt dependent shift of donor properties can be excluded. Error bars denote 95 % confident intervals.

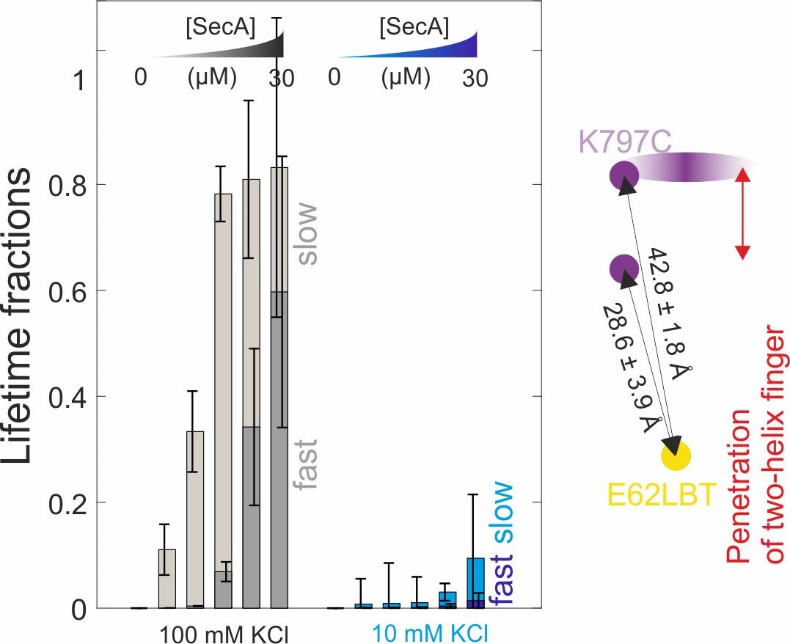

**Figure S6.** **Dependency of fluorescence lifetime partitioning between fast and slow fraction on SecA concentration.** Same as Fig. 3C but determined for another LRET pair (K797C on SecA and E62LBT on SecYEG; Average of n=2 independent experiments, respectively). Similar as for the G297LBT mutant, two distinct SecA-SecYEG complexes differing in the penetration depth of SecA’s two-helix finger into the SecY channel were observed. Calculated donor-acceptor distances represent an average of the pooled distances calculated from the respective 10 and 100 mM titrations (i.e. n=4).

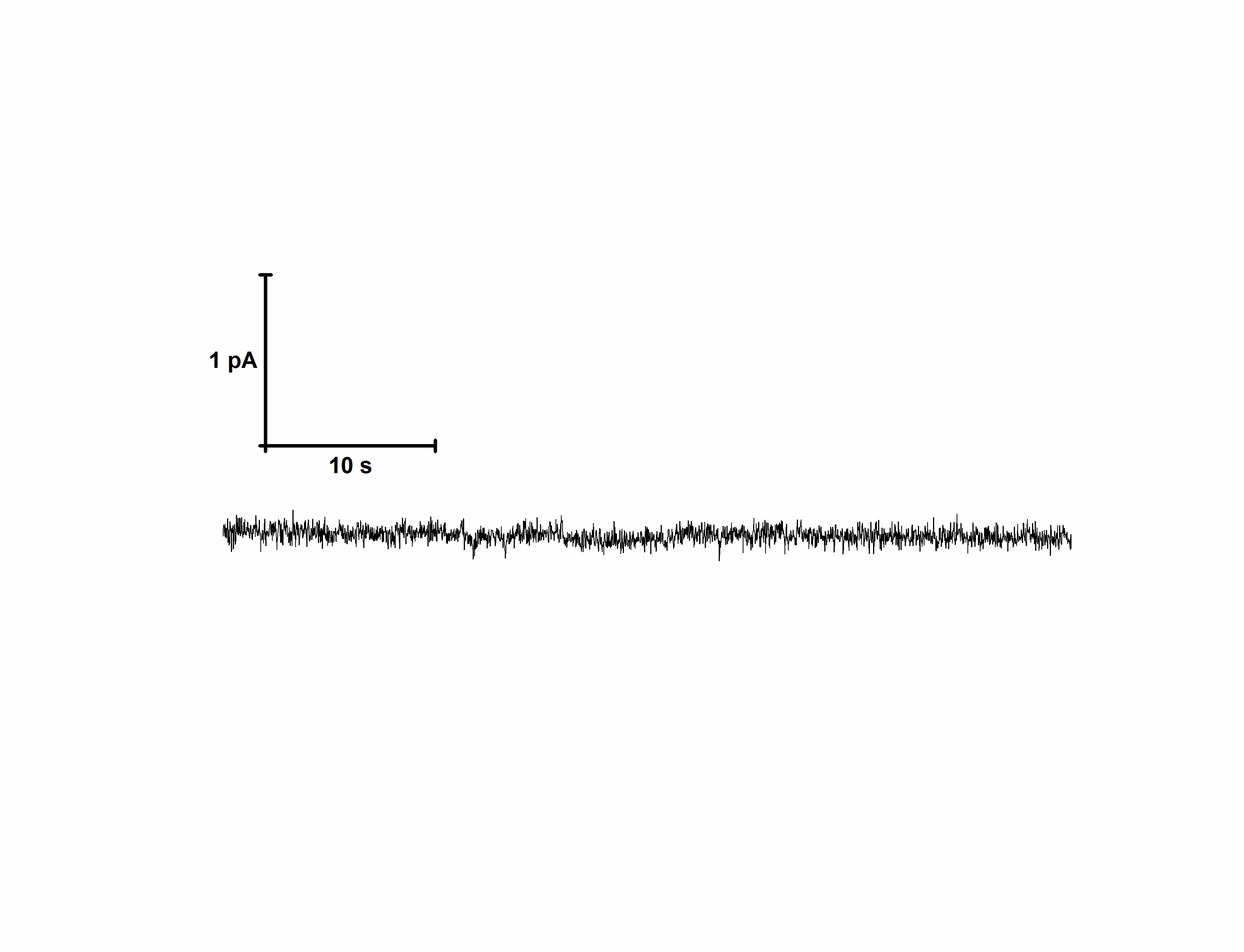

**Figure S7. No channel activity was observed in the absence of SecA.** Same conditions as in Fig. 4.

**Movie Legends**

**Movie S1.** HS-AFM movie of SecYEG prior to SecA injection. Last two frames mark the injection of SecA into the HS-AFM liquid cell. Image size: 200x200 nm², z-scale: 0-5.5 nm, scan speed: 400 ms/frame.

**Movie S2.** HS-AFM movie of SecA binding to SecYEG at low SecA surface concentration (continuation of Movie S1 after injection of SecA). Image size: 200x200 nm², z-scale: 0-5.5 nm, scan speed: 400 ms/frame.

**Movie S3.** HS-AFM movie of SecA binding to SecYEG at low SecA surface concentration. Image size: 200x200 nm², z-scale: 0-5.4 nm, scan speed: 472 ms/frame.

**Movie S2.** HS-AFM movie of SecA binding to SecYEG at low SecA surface concentration. Image size: 100x100 nm², z-scale: 0-5.3 nm, scan speed: 183 ms/frame.

**Movie S5.** HS-AFM movie of SecA binding to SecYEG at high SecA surface concentration. Image size: 200x200 nm², z-scale: 0-5.5 nm, scan speed: 100 ms/frame.
